## Supplementary data for "Genetic Determinants of pOXA-48 Plasmid Maintenance and Propagation in *Escherichia coli*"

**This PDF file includes:**

Figures S1 to S5
Table S1 to S4
SI References

**Other supporting materials for this manuscript include the following:**

Movie S1

**Figure S1**

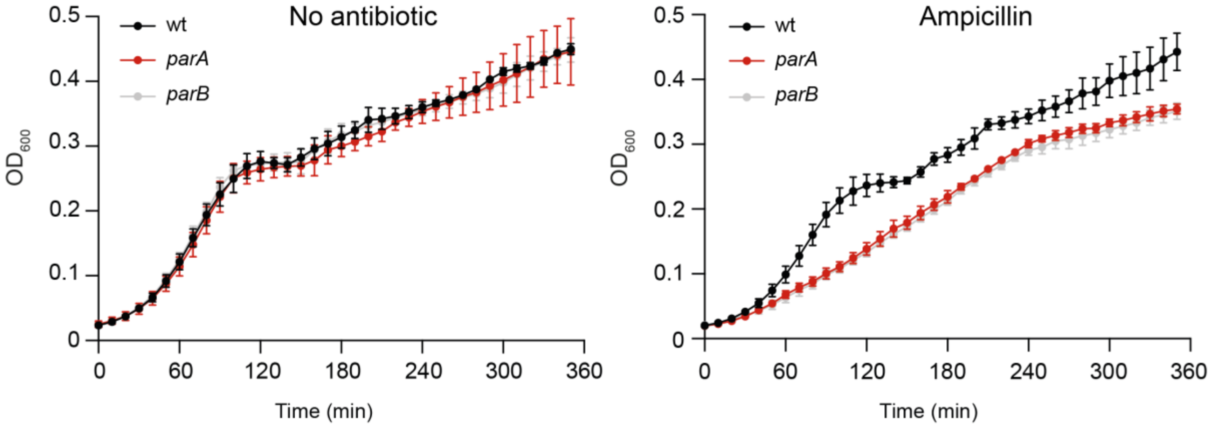


**Figure S1. Impact of selective pressure on *parA* and *parB* mutant growth.** Growth curve of wild-type (wt), *parA* and *parB* mutant strains represented as optical density (OD) at 600 nm wavelength measured during 360 min at 37°C in absence or presence of ampicillin. The mean and SD of three independent clones is represented for each strain.

**Figure S2**

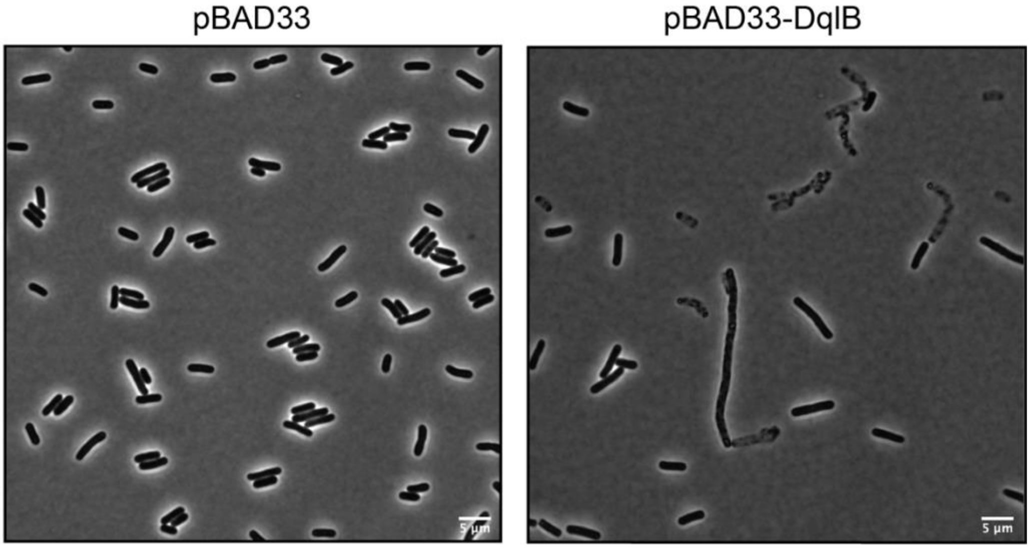


**Figure S2. Effect of DqlB production on cell morphology.** Representative phase contrast microscopy images of *E. coli* cells carrying either the empty vector pBAD33 (left; LY4076) or pBAD33-DqlB (right; LY4077), with the *dqlB* gene under the arabinose inducible $P_{BAD}$ promoter, after 4 h of induction with arabinose. Scale bar: 5 µm.

**Figure S3**

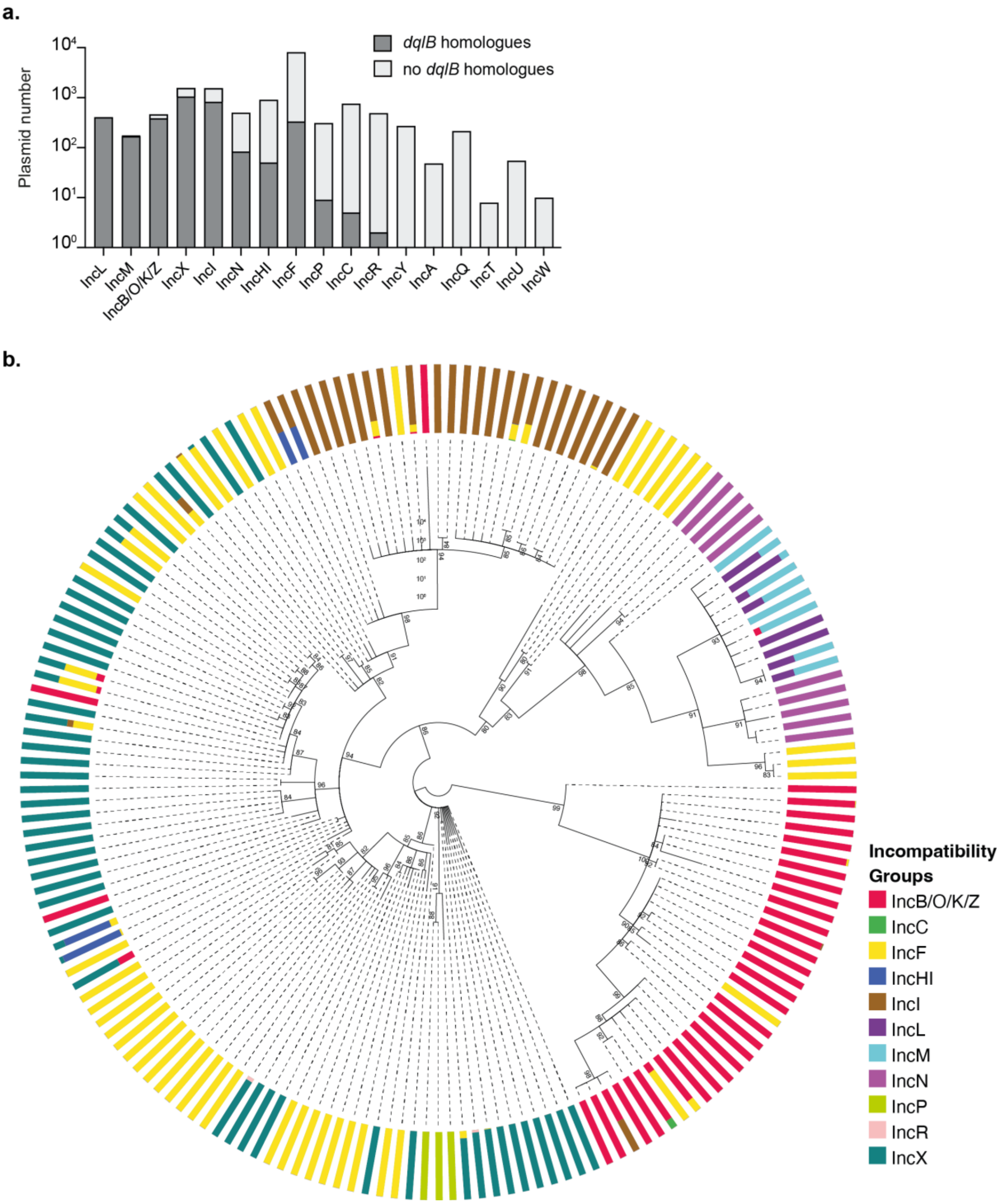


**Figure S3. Phylogenetic analysis and distribution of *dqlB* homologues among conjugative plasmids. a.** Distribution of *dqlB* homologues across different plasmid incompatibility groups. The number of plasmids within each incompatibility group is shown on the y-axis, with the presence (dark grey) or absence (light grey) of *dqlB* homologues indicated. **b**. Maximum likelihood phylogenetic tree of *dqlB* homologues identified from conjugative plasmid sequences. Branch support values were calculated using 1000 ultrafast bootstrap (UFboot). The outer ring color represents different incompatibility groups, demonstrating the phylogenetic relationships of *dqlB* homologues and their association with plasmid incompatibility groups.

**Figure S4**

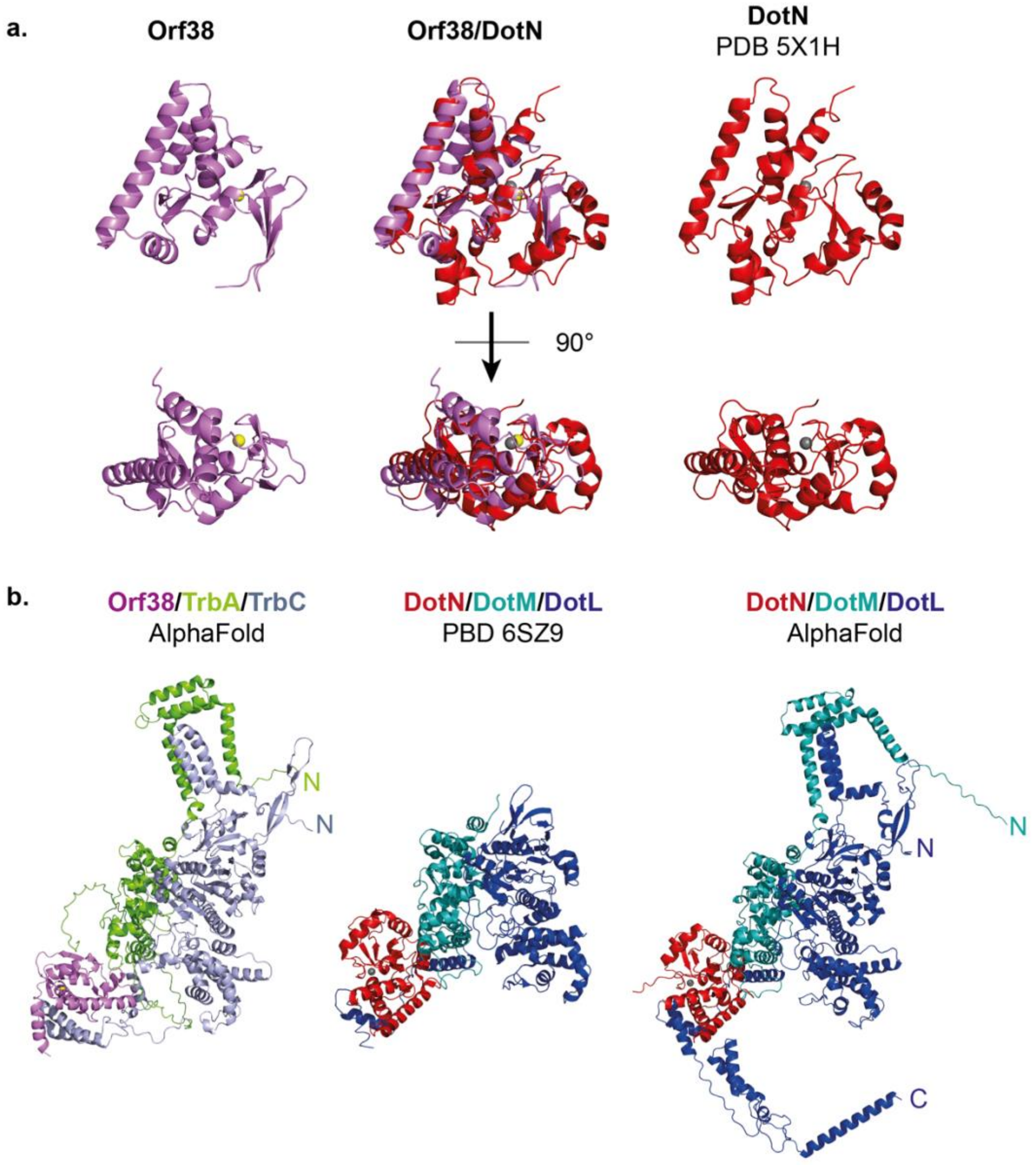


**Figure S4. Comparison of the Orf38 and DotN of *Legionella pneumophila* structures. a.** Left: predicted structure of Orf38 by AlphaFold 3.0; right: structure of DotN from *L. pneumophila* (PDB 5X1H); middle: structural alignment of the predicted Orf38 (purple) and DotN (red) using PyMoL. **b**. Comparison of the modeled Orf38/TrbA/TrbC complex with the DotN/DotM/DotL complex. Left: AlphaFold-predicted model of the Orf38/TrbA/TrbC complex, where Orf38 is shown in purple, TrbA in green, and TrbC in gray. Middle: Cryo-EM structure of the *L. pneumophila* DotN/DotM/DotL complex (PDB 6SZ9), with DotN in red, DotM in teal, and DotL in blue. Right: AlphaFold-predicted model of the DotN/DotM/DotL complex.

### Figure S5

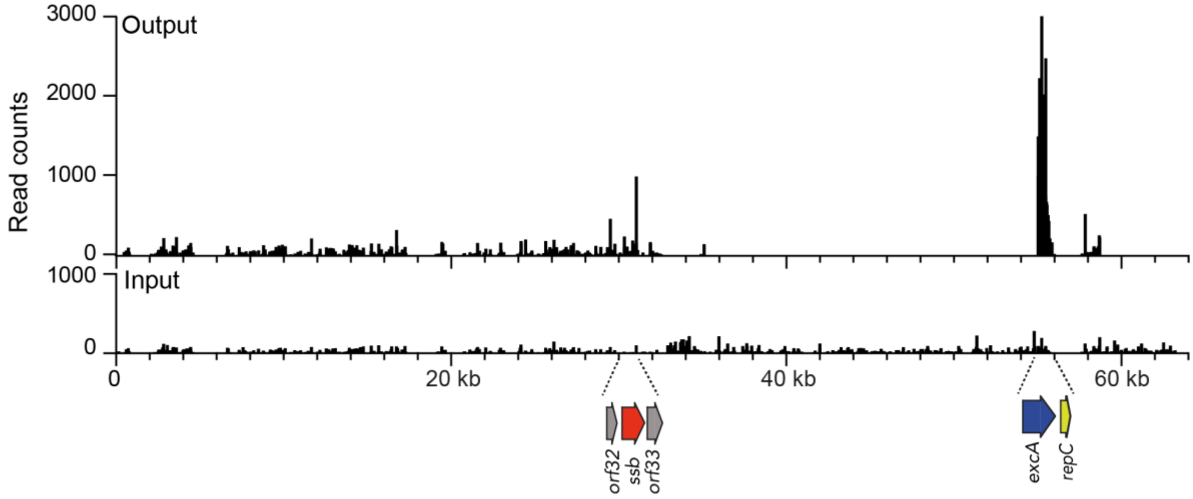


**Figure S5. Enrichment of transposon insertions in the output library compared to the input library.** Read distribution of transposon insertions across the pOXA-48 plasmid in both the input and output transposon libraries. The y-axis represents the number of reads, and the x-axis their position across the plasmid. Regions containing *orf32*, *ssb*, *orf33*, *excA*, and *repC*, with significant enrichment, are indicated.

**Table S1. Predicted proteins encoded by the pOXA-48 plasmid**

| Protein | Conserved domain | Putative function | Vir / Dot T4SS homologue | % identity pCTX-M3 |
|---|---|---|---|---|
| Orf67 | - | - | - | - |
| Tir* | CPBP family | Fertility inhibition | - | 93.5 |
| PemI | MazE superfamily | Type II antitoxin | - | 98.8 |
| PemK | CcdB superfamily | Type II toxin | - | 99 |
| MucA | LexA superfamily | Transcriptional repressor LexA | - | 98.6 |
| MucB | PolY superfamily | Translesion synthesis | - | 98.4 |
| Orf5 | - | - | - | 100 |
| Orf6 | - | - | - | 98.6 |
| Orf7 | - | - | - | 98.7 |
| Orf8 | HTH-XRE superfamily | Gene regulation | - | 94.4 |
| Orf9 | RelB Antitoxin | - | - | 90.6 |
| Orf10 | - | - | - | 89 |
| Orf10.1 | - | - | - | - |
| Orf11 | - | - | - | 90.2 |
| Orf13 | - | - | - | 72.9 |
| Orf14 | - | - | - | 96.7 |
| Orf15 | HTH-XRE superfamily | Gene regulation | - | 98.8 |
| Orf16 | Restriction nuclease | Host defense | - | 99 |
| Orf17 | - | - | - | 93.2 |
| Orf18 | - | - | - | 95.1 |
| ResD | XerD superfamily | Dimer resolution | - | 98.4 |
| ParA | StbA/ParM like | Partition | - | 99.7 |
| ParB | Plasmid_stab_B superfamily | Partition | - | 98.6 |
| Nuc | PLDc_Nuc | Nuclease | - | 96.2 |
| Orf19 | - | - | - | 99.4 |
| Orf20 | YiaG- HTH-XRE superfamily | Gene regulation | - | 99.4 |
| Orf21 | - | - | - | 97.8 |
| Orf22 | - | - | - | 98.8 |
| Orf23 | DUF308 | - | - | 99.2 |
| Orf24 | RadC superfamily | - | - | 98.7 |
| KorC | HTH-XRE superfamily | Gene regulation | - | 100 |
| Orf25* | DndB superfamily | - | - | 100 |
| CcgA1 | - | - | - | 91.5 |
| Orf26 | DUF4942 | - | - | 96.5 |
| Orf27 | - | - | - | 98.2 |
| RmoA | HHA superfamily | Gene regulation | - | 98.7 |
| Orf27.1 | - | - | - | 98.3 |
| Orf28 | - | - | - | 96.1 |
| Orf29 | DUF1380 | - | - | 98.5 |
| Orf30 | - | - | - | 99 |
| KlcA | Antirestriction superfamily | Host defense | - | 95.2 |
| Orf31 | - | - | - | 98.9 |
| Orf32 | - | - | - | 98.6 |
| Ssb | RPA_2b-aaRSs_OBF_like superfamily | Single strand binding protein | - | 97.9 |
| Orf33 | - | - | - | 99 |
| Orf34 | - | - | - | 99.4 |
| Orf35 | - | - | - | 97.5 |
| NikA | MobA superfamily | Reloxosome complex | - | 100 |
| NikB | PRK13878 superfamily | Relaxase | VirD2 | 99.7 |
| TraH | DotD superfamily | Outer membrane complex | DotD | 100 |
| TraI | TraI superfamily | Outer membrane lipoprotein | VirB7 / DotC | 100 |
| TraJ | Plasmid_TraJ superfamily | ATPase | VirB11 / DotB | 100 |
| TraK | IcmT/TraK | ATPase | IcmT | 100 |
| Pri | COG4643/LPD7 superfamily | Primase | - | 91.2 |
| TraL | - | - | - | 100 |
| Orf36.1 | Hha superfamily | Gene regulation | - | 100 |
| Orf36.2 | HNS_MvaT superfamily | Gene regulation | - | 100 |
| TraM | IcmL/DotI superfamily | Inner membrane complex | VirB8 / DotI | 99.6 |
| TraN | IcmK/DotH superfamily | Outer membrane complex | VirB9 / DotH | 99.7 |
| TraO | TraO_IncI1 superfamily | Cell envelope-spanning channel | VirB10 / DotG | 98.4 |

| | | | | |
|---|---|---|---|---|
| TraP | TraP_IncI1 superfamily | Pilus assembly | VirB3 / DotF | 94.9 |
| TraQ | TraQ_IncI1 superfamily | - | - | 99.4 |
| TraR | DUF6750 | - | - | 93 |
| Orf38 | - | - | - | 99.4 |
| TraU | VirB4 superfamily | ATPase | VirB4 / DotO | 99.2 |
| TraW | TraW superfamily | - | - | 100 |
| TraX | - | - | - | 53.7 |
| TraY | DotA/TraY superfamily | | DotA | 78.7 |
| ExcA | - | Entry exclusion | - | 34 |
| RepC | RepC_IncL/M superfamily | Plasmid copy number | - | 100 |
| RepB | Leader_RepB_LM superfamily | Replication | - | 87.5 |
| RepA | IncFII_repA superfamily | Replication | - | 99.4 |
| TrbC | TrbC superfamily | Type IV Coupling protein | VirD4 / DotL | 99.4 |
| TrbB | Thioredoxin_like superfamily | - | - | 98.7 |
| TrbA | DotM superfamily | - | DotM | 99.3 |
| TrbN | Lys-like superfamily | Lytic transglycosylase | - | 97.7 |

Elements of the Tn*1999* transposon and IS are not present. *gene encoding the protein is interrupted. Proteins that have been identified as being involved in plasmid maintenance and transfer are highlighted in yellow and blue, respectively.

**Table S2. Strain list.**

| Name | Genotype | Source or construct |
|---|---|---|
| MS388 | *E. coli* K-12 MG1655 *rpsL* (St$^R$) | Gift from F. Cornet |
| DY330 | *W3110 ΔlacU169, gal490, λcI857, Δ(cro-bioA)* | [1] |
| DH5α | F$^-$ *endA1 glnV44 thi-1 recA1 relA1 gyrA96 deoR nupG purB20* φ80d*lacZ*ΔM15 Δ(*lacZYA-argF*)U169, hsdR17($r_K^-m_K^+$), λ | Lab collection |
| LY1898 | MFDpir / pSAM_Ec (Ap$^R$, Kan$^R$, DAP-) | [2] |
| LY945 | MS388 *ilvA::Cm* (St$^R$, Cm$^R$) | [3] |
| LY1413 | *E. coli* Top10 / pOXA-48 (Ap$^R$) | [4] |
| LY1507 | MS388 *ilvA::erm* / pOXA-48 (St$^R$, Erm$^R$, Ap$^R$) | LY1413 x LY832 to Erm$^R$ Ap$^R$ |
| LY1672 | DY330 / pOXA-48 (Ap$^R$) | LY1507 x DY330 to Ap$^R$ lac- |
| LY1844 | MS388 / pOXA-48 (St$^R$, Ap$^R$) | LY1672 x MS388 to Ap$^R$ St$^R$ |
| LY3154 | DY330 / pOXA-48-*ΔparA::frt-kan-frt* (Ap$^R$, Kan$^R$) | λRed *frt-kan-frt* (OL1616/OL1617) at the endogenous pOXA-48 locus in LY1672 |
| LY3162 | MS388 / pOXA-48-*ΔparA::frt-kan-frt* (St$^R$, Ap$^R$, Kan$^R$) | Conjugation LY3154 x MS388 to St$^R$ Kan$^R$ |
| LY3257 | MS388 / pOXA-48-*ΔparA::frt* (St$^R$, Ap$^R$) | Derivative of LY3162, *kan* removed via pCP20 |
| LY3155 | DY330 / pOXA-48-*ΔparB::frt-kan-frt* (Ap$^R$, Kan$^R$) | λRed *frt-kan-frt* (OL1620/OL1621) at the endogenous pOXA-48 locus in LY1672 |
| LY3163 | MS388 / pOXA-48-*ΔparB::frt-kan-frt* (St$^R$, Ap$^R$, Kan$^R$) | Conjugation LY3155 x MS388 to St$^R$ Kan$^R$ |
| LY3352 | MS388 / pOXA-48-*ΔparB::frt* (St$^R$, Ap$^R$) | Derivative of LY3163, *kan* removed via pCP20 |
| LY3156 | DY330 / pOXA-48-*Δnuc::frt-kan-frt* (Ap$^R$, Kan$^R$) | λRed frt-kan-frt (OL1622/OL1623) at the endogenous pOXA-48 locus in LY1672 |
| LY3164 | MS388 / pOXA-48-*Δnuc::frt-kan-frt* (St$^R$, Ap$^R$, Kan$^R$) | Conjugation LY3156 x MS388 to St$^R$ Kan$^R$ |
| LY3258 | MS388 / pOXA-48-*Δnuc::frt* (St$^R$, Ap$^R$) | Derivative of LY3164, *kan* removed via pCP20 |
| LY4049 | DY330 / pOXA-48-*Δorf19::frt-kan-frt* (Ap$^R$, Kan$^R$) | λRed *frt-kan-frt* (OL2218/OL2219) at the endogenous pOXA-48 locus in LY1672 |
| LY832 | MS388 *ilvA::erm* (St$^R$, Erm$^R$) | [5] |
| LY4141 | MS388 *ilvA::erm* / pOXA-48-*Δorf19::frt-kan-frt* (St$^R$, Erm$^R$, Ap$^R$, Kan$^R$) | Conjugation LY4049 x LY832 to Erm$^R$Kn$^R$ |
| LY2172 | DY330 / pOXA-48-P$_{Tet}$-*sfgfp*-frt-kn-frt (Ap$^R$, Kan$^R$) | λRed *frt-kan-frt* (OL974/OL975) at the *orf27*.1 pOXA-48 locus in LY1672 |
| LY4146 | DY330 / pOXA-48-P$_{Tet}$-*sfgfp*-frt-kn-frt-*Δorf20::frt-cat-frt* (Ap$^R$, Kan$^R$, Cm$^R$) | λRed *frt-cat-frt* (OL2220/OL2221) at the endogenous locus in LY2172 |
| LY681 | MS388 *ilvA*::P$_{lacIQ1}$-tetR-frt-kan-frt (St$^R$) | [3] |

| | | |
|---|---|---|
| LY699 | MS388 *ilvA::*$P_{lacIQ1}$-tetR ($St^R$) | Derivative of LY681, *kan* removed via pCP20 |
| LY4138 | MS388 *ilvA::*$P_{lacIQ1}$-tetR / pOrf20 ($St^R$, $Gm^R$) | Transformation LY699 with pOrf20 |
| LY4161 | MS388 *ilvA::*$P_{lacIQ1}$-tetR / pOrf20 / pOXA-48-$P_{Tet}$-*sfgfp*-frt-kn-frt-*Δorf20::frt-kan-frt* ($St^R$, $Ap^R$, $Kan^R$, $Cm^R$, $Gm^R$) | Conjugation LY4146 x LY4138 to $Gm^R$ $Kan^R$ $Cm^R$ |
| LY1593 | MS388 *ilvA*::$P_{tac}$-mcherry ($St^R$, $Gm^R$) | [5] |
| LY4140 | MS388 *ilvA*::$P_{tac}$-mcherry / pOrf20($St^R$) | Transformation LY1593 with pOrf20 |
| LY4129 | DY330 / pOXA-48-*Δorf32::frt-kan-frt* ($Ap^R$, $Kan^R$) | λRed *frt-kan-frt* (OL2284/OL2285) at the endogenous pOXA-48 locus in LY1672 |
| LY4132 | MS388 / pOXA-48-*Δorf32::frt-kan-frt* ($St^R$, $Ap^R$, $Kan^R$) | Conjugation LY4129 x MS388 to $St^R$ $Kan^R$ |
| LY4148 | MS388 / pOXA-48-*Δorf32::frt* ($St^R$, $Ap^R$) | Derivative of LY4132, *kan* removed via pCP20 |
| LY4192 | MS388 / pOXA-48-*Δorf32::frt* / pOrf32 ($St^R$, $Ap^R$, $Gm^R$) | Transformation LY4148 with pOrf32 |
| LY4130 | DY330 / pOXA-48-*Δssb::frt-kan-frt* ($Ap^R$, $Kan^R$) | λRed *frt-kan-frt* (OL2286/OL2287) at the endogenous pOXA-48 locus in LY1672 |
| LY4133 | MS388 / pOXA-48-*Δssb::frt-kan-frt* ($St^R$, $Ap^R$, $Kan^R$) | Conjugation LY4130 x MS388 to $St^R$ $Kan^R$ |
| LY4149 | MS388 / pOXA-48-*Δssb::frt* ($St^R$, $Ap^R$) | Derivative of LY4133, *kan* removed via pCP20 |
| LY4151 | DY330 / pOXA-48-*Δorf33::frt-kan-frt* ($Ap^R$, $Kan^R$) | λRed *frt-kan-frt* (OL2288/OL2289) at the endogenous pOXA-48 locus in LY1672 |
| LY4152 | MS388 / pOXA-48-*Δorf33::frt-kan-frt* ($St^R$, $Ap^R$, $Kan^R$) | Conjugation LY4151 x MS388 to $St^R$ $Kan^R$ |
| LY4153 | MS388 / pOXA-48-*Δorf33::frt* ($St^R$, $Ap^R$ | Derivative of LY4152, *kan* removed via pCP20 |
| LY3245 | DY330 / pOXA-48-*Δorf36.1::frt-kan-frt* ($Ap^R$, $Kan^R$) | λRed *frt-kan-frt* (OL1721/OL1722) at the endogenous pOXA-48 locus in LY1672 |
| LY3270 | MS388 / pOXA-48-*Δorf36.1::frt-kan-frt* ($St^R$, $Ap^R$, $Kan^R$) | Conjugation LY33245 x MS388 to $St^R$ $Kan^R$ |
| LY3303 | MS388 / pOXA-48-*Δorf36.1::frt* ($St^R$, $Ap^R$) | Derivative of LY3270, *kan* removed via pCP20 |
| LY3697 | MS388 / pOXA-48-*Δorf36.1::frt* /pOrf36.1 ($St^R$, $Ap^R$) | Transformation LY3303 with pOrf36.1c |
| LY3378 | DY330 / pOXA-48-*Δorf38::frt-kan-frt* ($Ap^R$, $Kan^R$) | λRed *frt-kan-frt* (OL1825/OL1826) at the endogenous pOXA-48 locus in LY1672 |
| LY3698 | DY330 / pOXA-48-*Δorf38::frt-kan-frt* /pOrf38 ($Ap^R$, $Kan^R$, $Gm^R$) | Transformation LY3378 with pOrf38 |
| LY3710 | MS388 / pOXA-48-*Δorf38::frt-kan-frt* ($St^R$, $Ap^R$, $Kan^R$) | Conjugation LY3698 x MS388 to $St^R$ $Kan^R$ |
| LY3739 | MS388 / pOXA-48-*Δorf38::frt* ($St^R$, $Ap^R$) | Derivative of LY3710, *kan* removed via pCP20 |
| LY3747 | MS388 / pOXA-48-*Δorf38::frt* /pOrf38($St^R$, $Ap^R$) | Transformation LY3739 with pOrf38 |
| LY3158 | DY330 / pOXA-48-*ΔexcA::frt-kan-frt* ($Ap^R$, $Kan^R$) | λRed *frt-kan-frt* (OL1624/OL1625) at the endogenous pOXA-48 locus in LY1672 |
| LY3166 | MS388 / pOXA-48-*ΔexcA::frt-kan-frt* ($St^R$, $Ap^R$, $Kan^R$) | Conjugation LY3158 x MS388 to $St^R$ $Kan^R$ |
| LY3260 | MS388 / pOXA-48-*ΔexcA::frt* ($St^R$, $Ap^R$) | Derivative of LY3166, *kan* removed via pCP20 |
| LY3608 | MS388 / pOXA-48-*ΔexcA::frt* / pExcA | Transformation LY3260 with pExcA |
| LY4164 | Tn insertion *excA* clone 1 | Tn-seq output library |
| LY4165 | Tn insertion *excA* clone 2 | Tn-seq output library |
| LY3157 | DY330 / pOXA-48-*ΔrepC::frt-kan-frt* ($Ap^R$, $Kan^R$) | λRed *frt-kan-frt* (OL1626/OL1627) at the endogenous pOXA-48 locus in LY1672 |
| LY3165 | MS388 / pOXA-48-*ΔrepC::frt-kan-frt* ($St^R$, $Ap^R$, $Kan^R$) | Conjugation LY3157 x MS388 to $St^R$ $Kan^R$ |
| LY3259 | MS388 / pOXA-48-*ΔrepC::frt* ($St^R$, $Ap^R$) | Derivative of LY3165, *kan* removed via pCP20 |
| LY3606 | MS388 / pOXA-48-*ΔrepC::frt* / pRepC | Transformation LY3259 with pRepC |
| LY1361 | TB28 *ilvA::IsceICS- frt-cat-frt* / F-Tn*10* ($Cm^{R,}$ $Tc^R$) | [5] |
| LY1369 | TB28 *ilvA::IsceICS- frt-cat-frt* / F-Tn*10* / pBG29 ($Cm^R$, $Tc^R$, $Kn^R$) | [5] |
| LY1522 | TB28 *ilvA::IsceICS- frt-cat-frt* / F-Tn*10* / pBG50 ($Cm^R$, $Tc^R$, $Kn^R$) | [5] |

| | | |
|---|---|---|
| LY1549 | TB28 *ilvA::IsceICS- frt-cat-frt* / F-Tn*10* / pBG52 ($Cm^R$, $Tc^R$, $Kn^R$) | [5] |
| LY4038 | TB28 *ilvA::IsceICS- frt-cat-frt* / F-Tn*10* / $TAP_{kn}$-Cas9-OXA48-argB ($Cm^R$, $Tc^R$, $Kn^R$) | Transformation of LY1361 with $TAP_{kn}$-OXA48-*agrB* |
| LY4076 | MS388 / pBAD33 ($St^R$, $Cm^R$) | Transformation MS388 with pBAD33 |
| LY4077 | MS388 / pBAD33-DqlB ($St^R$, $Cm^R$) | Transformation MS388 with pBAD33-DqlB |
| LY4078 | MS388 / pBG50 / pBAD33 ($St^R$, $Kn^{R,}$ $Cm^R$) | Transformation of MS388 with pBG50 and pBAD33 |
| LY4079 | MS388 / pBG50 / pBAD33-DqlB ($St^R$, $Kn^{R,}$ $Cm^R$) | Transformation of MS388 with pBG50 and pBAD33-DqlB |
| LY4080 | MS388 / $TAP_{kn}$-Cas9-OXA48-argB/ pBAD33 ($St^R$,Cm $^R$, $Kn^R$) | Transformation of MS388 with TAP-OXA48-agrB and pBAD33 |
| LY4081 | MS388 / $TAP_{kn}$-Cas9-OXA48-argB / pBAD33-DqlB ($St^R$,Cm $^R$, $Kn^R$) | Transformation of MS388 with TAP-OXA48-agrB and pBAD33-DqlB |

Abbreviation *cat* and *kan* refers to insertion conferring resistance to chloramphenicol and kanamycin. $St^R$, $Tc^R$ $Ap^R$, $Cm^R$, $Kan^R$ and $Erm^R$ refer to streptomycin, tetracycline, ampicillin, chloramphenicol, kanamycin and erythromycin resistances, respectively. *frt* refers to the FLP site-specific recombination site.

### Table S3: plasmid list.

| Name | Usage | Reference or construct |
|---|---|---|
| pOXA-48 | Carries the $bla_{OXA-48}$ gene that encodes the OXA48 carbapenemase | [4] |
| F-Tn*10* | Template for $P_{Tet}$ promoter | [3] |
| pSAM_Ec | Carries the Himar-1 transposase used for Tn-Seq library, $Ap^R$, $Kan^R$ | [6] |
| pKD3 | Template plasmid for *frt*-flanked *cat* cassette, $Cm^R$ | [7] |
| pR6K-sfGFP | Template plasmid for *frt*-flanked *kan* cassette, $Kan^R$ | [3] |
| pCP20 | Flp expression plasmid, $Ap^R$, $Cm^R$, ts | [7] |
| pUC18T-mini-Tn7T-Gm-Dsredexpress | Carries the *PA1/04/03* promoter, $Ap^R$, $Gm^R$ | [8] |
| pUA66-psulA-gfp-Gm | Carries the pSC101 *oriV* and *acc3* gene, $Gm^R$ | Derivative of pUA66[9] |
| pTrc99a | Carries the *lacI* gene, $Ap^R$ | GE healthcare |
| pSC101-pPA | Expression vector used for complementation experiments. | Assembly of pSC101 *oriV* - *acc3* gene from pUA66-psulA-gfp-Gm with *PA1/04/03* promoter from pUC18T-mini-Tn7T-Gm-Dsredexpress and *lacI* gene from pTrc99a. |
| pR6K-ptet-sfGFP | Carries the *sfgfp* gene under the $P_{Tet}$ promoter, $Kan^R$ | Insertion of $P_{Tet}$ from F-Tn*10* upstream of *sfgfp* pR6K-sfGFP |
| pOrf20 | Carries *orf20* gene under the *PA1/04/03* promoter, $Gm^R$ | Insertion of *orf20* from pOXA-48 in pSC101-pPA |
| pOrf36.1 (pBG77) | Carries *orf36.1* gene under the *PA1/04/03* promoter, $Gm^R$ | Insertion of *orf36.1* from pOXA-48 in pSC101-pPA |
| pOrf38 (pBG79) | Carries *orf38* gene under the *PA1/04/03* promoter, $Gm^R$ | Insertion of *orf38* from pOXA-48 in pSC101-pPA |
| pExcA | Carries *excA* gene he *PA1/04/03* promoter, $Gm^R$ | Insertion of *excA* from pOXA-48 in pSC101-pPA |
| pRepC | Carries *repC* gene under the *PA1/04/03* promoter, $Gm^R$ | Insertion of *repC* from pOXA-48 in pSC101-pPA |
| pOrf32 | Carries *orf32* gene under the *PA1/04/03* promoter, $Gm^R$ | Insertion of *orf32* from pOXA-48 in pSC101-pPA |
| $TAP_{kn}$-Cas9-nsp (pBG29) | Produces Cas9 and nsp spacer, $Kn^R$ | [5] |
| $TAP_{kn}$-Cas9-OXA48 (pBG50) | Produces Cas9 with OXA48 spacer targeting the promoter region of $bla_{OXA48}$, $Kn^R$ | [5] |

| | | |
|---|---|---|
| $TAP_{kn}$-Cas9-OXA48-pemI (pBG52) | Produces PemI and Cas9 with OXA48 spacer targeting the promoter region of *bla*$_{OXA48}$, $Kn^R$ | [5] |
| $TAP_{kn}$-Cas9-OXA48-pemI-agrB | Produces PemI and *argB* and Cas9 with OXA48 spacer targeting the promoter region of *bla*$_{OXA48}$, $Kn^R$. | Insertion *argB* under its own promoter from pOXA-48 in pBG52 |
| $TAP_{kn}$-Cas9-OXA48-agrB | Produces *argB* and Cas9 with OXA48 spacer targeting the promoter region of *bla*$_{OXA48}$, $Kn^R$. | Deletion *pemI* in $TAP_{kn}$-Cas9-OXA48-pemI-agrB |
| pBAD33-DqlB | Carries *dqlB* gene under the $P_{BAD}$ promoter. | Insertion *dqlB* from pOXA-48 in pBAD33 plasmid |

Abbreviations: *acc3*, *cat* and *kan* refers to gene conferring resistance to gentamycin, chloramphenicol and kanamycin. $Tc^R$, $Amp^R$, $Cm^R$, $Gm^R$ and $Kan^R$ refer to tetracycline, ampicillin, chloramphenicol, gentamycin and kanamycin resistances respectively. *frt* and ts refer to the FLP site-specific recombination site and temperature sensitive.

### Table S4: primer list.

| Primer | Sequence | Construct |
|---|---|---|
| OL1306 | ATGCAAGCTCTAGAGGCATC | pSC101-pPA construction by Gibson Assembly (GA) cloning |
| OL680 | GGCTGACGCCGTTGGATACACCAAG | |
| OL1691 | CTTGGTGTATCCAACGGCGTCAGCCAATTCGCGCGCGAAGGCGAA | |
| OL1692 | GGTGTCAACGTCACTGCCCGCTTTCTCAGAATATTTGCCAGAACC | |
| OL713 | GAAAGCGGGCAGTGACGTTGACACCCAAGGGAAAATTTATCAAAA | |
| OL1693 | TATTTGATGCCTCTAGAGCTTGCATGCTTAATTTCTCCTCTTTAA | |
| OL2226 | GGAACGACGCTAACCATCTTAATCATGCTTAATTTCTCCTCTT | pOrf20 construction by GA cloning |
| OL2227 | CAGCTGGTTGAGAAGGCGAAATAACAAGCTCTAGAGGCATCAAA | |
| OL2228 | ATTAAGATGGTTAGCGTCGTTCCG | |
| OL2229 | TTATTTCGCCTTCTCAACCAGCTG | |
| OL1892 | CATGCTTAATTTCTCCTCTT | pOrf36.1 construction by GA cloning |
| OL1893 | CAAGCTCTAGAGGCATCAAA | |
| OL1894 | AATTAAAGAGGAGAAATTAAGCATGAAATCAGCTGAAGACTG | |
| OL1895 | TTTTATTTGATGCCTCTAGAGCTTGCTACCGGAAGAGTCCTTTCG | |
| OL1892 | CATGCTTAATTTCTCCTCTT | pOrf38 construction by GA cloning |
| OL1893 | CAAGCTCTAGAGGCATCAAA | |
| OL1900 | AATTAAAGAGGAGAAATTAAGCATGAAAGATTTTAAGCTCAC | |
| OL1901 | TTTTATTTGATGCCTCTAGAGCTTGTCATTTAAAACCTCTCTTAC | |
| OL1878 | CTAGAGTCGACCTGCAGGCATGCAATGCAAGCTCTAGAGGCATC | pExcA construction by GA cloning |
| OL1879 | GAGCTCGAATTCGCTAGCCCAAAAATTCTAGATGTGTGAAATT | |
| OL1636 | TTTTTGGGCTAGCGAATTCGAGCTCTGGTATAGCGGGGGCGTGAT | |
| OL1637 | TGCATGCCTGCAGGTCGACTCTAGATCAATACGTTCGTGGTAGCA | |
| OL1878 | CTAGAGTCGACCTGCAGGCATGCAATGCAAGCTCTAGAGGCATC | pRepC construction by GA cloning |
| OL1879 | GAGCTCGAATTCGCTAGCCCAAAAATTCTAGATGTGTGAAATT | |
| OL1638 | TTTTTGGGCTAGCGAATTCGAGCTCCGTATGCTGTGGGAGGAAGA | |
| OL1639 | TGCATGCCTGCAGGTCGACTCTAGATCACTCTTCAGAATCGTCAA | |
| OL2326 | AAAGAAGCAGACCAAAAAAAAGTGAATGCCG | pOrf32 construction by GA cloning |
| OL2327 | TCAGTAAAACTTCACGTTCGATTTCAGGG | |
| OL2328 | CGGCATTCACTTTTTTTTGGTCTGCTTCTTTCATGCTTAATTTCTCCT CTT | |
| OL2329 | GCCCTGAAATCGAACGTGAAGTTTTACTGACAAGCTCTAGAGGCAT CAAA | |
| OL374 | ATGTCTAAAGGTGAAGAACTGTTC | pR6K-$P_{Tet}$-sfGFP construction by GA cloning |
| OL375 | CACTGCCCGCTTTCCAGT | |
| OL871 | CTGGAAAGCGGGCAGTGTAATTCCTAATTTTTGTTGACAC | |
| OL872 | GTTCTTCACCTTTAGACATTTCACTTTTCTCTATCACTGA | |
| OL1616 | TGATTTCTACGGTTTTTTTCTAAATAATGTGAGGTTCAAAGTGTAG GCTGGAGCTGCTTC | λred *parA* deletion |
| OL1617 | GCACTTACGGTTATCAGTCATACTCAGCCCTTCATTGCCACATATG AATATCCTCCTTAG | |

| | | |
|---|---|---|
| OL1620 | TCGATCTGGTTCTGGGTATGGTGGCAATGAAGGGCTGAGTGTGTAG GCTGGAGCTGCTTC | $\lambda$red *parB* deletion |
| OL1621 | AACAGCACATATTTTTTCACGCAGCCCCCTTAATCATCATCATATG AATATCCTCCTTAG | |
| OL1622 | CGTCCAGCATCTGTTCCCGGATGATGATTAAGGGGGCTGCGGTGTA GGCTGGAGCTGCTT | $\lambda$red *nuc* deletion |
| OL1623 | GTTAAAAATCAGGCGGCAGAGTTTCTGCCGCTGCTGTAGGCATATG AATATCCTCCTTAG | |
| OL2218 | CATTTTTCGGTAGCGCCCCGACTGGCGAAGGAGTAAAACCGTGTA GGCTGGAGCTGCTTC | $\lambda$red *orf19* deletion |
| OL2219 | CATCTTAATCATTACCCTTCTCCCAGAAGTCTGAATTCAGCATATG AATATCCTCCTTAG | |
| OL2220 | GTGAAAATTTTTCTGAATTCAGACTTCTGGGAGAAGGGTAGTGTAG GCTGGAGCTGCTTC | $\lambda$red *orf20* deletion |
| OL2221 | TGTGGTGATCATAAAGGGTCCTCTCTGGTCGCCGCGGGCCCATATG AATATCCTCCTTAG | |
| OL974 | TCAGAAGAGGTCATAATGCACCTGTAAGGCTGTTTAGGAACTGGC ACGACAGGTTTCCCGA | Insertion $P_{Tet}$-*sfgfp-frt-kan-frt* in pOXA-48 |
| OL975 | CATTAGACGTATAATGTTTTTACTGATTCAAATCAGTACCCATATG AATATCCTCCTTAGT | |
| OL1721 | TTTCAACGCGGCGGTGTTCCGCCGCGGTTAGTAAACGCATGTGTAG GCTGGAGCTGCTTC | $\lambda$red *orf36.1* deletion |
| OL1722 | AGACTGGCTGCATACTGTCAGACGTTTCATGAATGAAGATCATATG AATATCCTCCTTAG | |
| OL1825 | AATACCGGAGTTCCCCCATGAAAGATTTTAAGCTCACCAGGTGTAG GCTGGAGCTGCTTC | $\lambda$red *orf38*deletion |
| OL1826 | CATCCTCAAAGAAAGAAATCATTTAAAACCTCTCTTACCCCATATG AATATCCTCCTTAG | |
| OL2284 | AAAAAATGGGAGGATTAAGGGGGATTTCTCCCCCTCTTTTCCATAT GAATATCCTCCTTAG | $\lambda$red *orf32* deletion |
| OL2285 | AGTCTTTCGCGGTAGCGCCCCGACTGGCGAAGGAGTTGAAGTGTA GGCTGGAGCTGCTTC | |
| OL2286 | CCATTTTTTACTGATAAACATCAAATATATGAGGTGTATTGTGTAG GCTGGAGCTGCTTC | $\lambda$red *ssb* deletion |
| OL2287 | TTGGCGGCCCGCGTCCTGCGGGCCGTTCTAACTCAAATACCCATAT GAATATCCTCCTTAG | |
| OL2288 | AGAAAAAAATGCCCCTTTTCGAGGCAAATTTTTAACCGGTCCATAT GAATATCCTCCTTAG | $\lambda$red *orf33* deletion |
| OL2289 | CGGCCCGCAGGACGCGGGCCGCCAATCCAGAGGGGGGAGTGTGTA GGCTGGAGCTGCTTC | |
| OL1624 | ACGGTTGATAAAGACAAAGATGGTATAGCGGGGGCGTGATGTGTA GGCTGGAGCTGCTTC | $\lambda$red *excA* deletion |
| OL1625 | CATGAAAAAACCGCCTTTTCAGGCGGTTTGGAAGTAAACACATAT GAATATCCTCCTTAG | |
| OL1626 | TATACTCAGGTGGTTATAGTCGTATGCTGTGGGAGGAAGAGTGTAG GCTGGAGCTGCTTC | $\lambda$red *repC* deletion |
| OL1627 | AGCGGGGCAATACCTGCGAACCCGCTTTTACATAGTAAGACATAT GAATATCCTCCTTAG | |
| OL2209 | CTCGAGCGCTCGGTACCCACAGAGAGCGTTTCAGGC | $TAP_{kn}$-Cas9-OXA48-pemI-argB construction by insertion in SmaI-digested pBG52 |
| OL2210 | GCACTGAAGGATCCCCCCGTATACCCTATTCCCGGTGG | |
| OL602 | GATCCTCTAGAGTCGACCTGCAGGC | $TAP_{kn}$-Cas9-OXA48-argB construction by removing *pemI*, followed by phosphorylation and self-ligation |
| OL1131 | GAATGACCTGTCACGGGATAATCCGGCGGCAAACAGCAAAGACGT | |
| OL2245 | GTTTTGGCGGATGAGAGAAG | pBAD-DqlB construction by GA cloning |
| OL2246 | ATGGAGAAACAGTAGAGAGTTGCG | |
| OL2247 | CTCTCTACTGTTTCTCCATGTCAATCTGTTGACGTTCTTC | |
| OL2248 | CTTCTCTCATCCGCCAAAACCGATGATCGAGATGTAACCC | |
| L1204-I1 | CAAGCAGAAGACGGCATACGAGATCGTGATAGACCGGGGACTTAT CATCCAACCTGT | Tn-seq libraries |
| L1204-I2 | CAAGCAGAAGACGGCATACGAGATGCCTAAAGACCGGGGACTTAT CATCCAACCTGT | |

**Movie S1 (separate file). Transconjugant viability following acquisition of the *orf20* mutant plasmid under selective pressure.** Donor and recipient cells were mixed for 4 hours on a filter before being imaged every 10 minutes by time-lapse microscopy, with a continuous flow of fresh medium supplemented with ampicillin. Recipient cells, exhibiting red mCherry fluorescence, are sensitive to ampicillin. In the absence of complementation of the *orf20* mutation by the pOrf20 plasmid (R without pOrf20, right panel), some recipient cells appear elongated with an increased width, continuing to elongate over time before undergoing bulging and eventual cell lysis. Non-fluorescent donor cells and transconjugants (which exhibit both mCherry and sfGFP fluorescence) are resistant to ampicillin due to expression of the $bla_{\mathrm{OXA\text{-}48}}$ gene encoded by pOXA-48. Donors *E. coli ilvA*::$P_{\mathrm{Biofab}}$-*tetR* / pOXA-48-Δorf20-$P_{\mathrm{tetR}}$-sfGFP/ pOrf20 (LY4161); recipients left panel: *E. coli ilvA*::$P_{\mathrm{tac}}$-*mcherry* / pOrf20 (LY4140), right panel *E. coli ilvA*::$P_{\mathrm{tac}}$-*mcherry* (LY1593). Scale bar: 10 μm.

### SI References


1. Yu, D. *et al.* An efficient recombination system for chromosome engineering in Escherichia coli. *Proc. Natl. Acad. Sci. U.S.A.* **97**, 5978–5983 (2000).
2. Royet, K., Parisot, N., Rodrigue, A., Gueguen, E. & Condemine, G. Identification by Tn-seq of Dickeya dadantii genes required for survival in chicory plants. *Mol Plant Pathol* **20**, 287–306 (2019).
3. Nolivos, S. *et al.* Role of AcrAB-TolC multidrug efflux pump in drug-resistance acquisition by plasmid transfer. *Science* **364**, 778–782 (2019).
4. Poirel, L., Bonnin, R. A. & Nordmann, P. Genetic Features of the Widespread Plasmid Coding for the Carbapenemase OXA-48. *Antimicrobial Agents and Chemotherapy* **56**, 559–562 (2012).
5. Reuter, A. *et al.* Targeted-antibacterial-plasmids (TAPs) combining conjugation and CRISPR/Cas systems achieve strain-specific antibacterial activity. *Nucleic Acids Res* **49**, 3584–3598 (2021).
6. Goodman, A. L. *et al.* Identifying genetic determinants needed to establish a human gut symbiont in its habitat. *Cell Host Microbe* **6**, 279–289 (2009).
7. Datsenko, K. A. & Wanner, B. L. One-step inactivation of chromosomal genes in Escherichia coli K-12 using PCR products. *Proc. Natl. Acad. Sci. U.S.A.* **97**, 6640–6645 (2000).
8. Choi, K.-H. & Schweizer, H. P. mini-Tn7 insertion in bacteria with single attTn7 sites: example Pseudomonas aeruginosa. *Nat Protoc* **1**, 153–161 (2006).
9. Zaslaver, A. *et al.* A comprehensive library of fluorescent transcriptional reporters for Escherichia coli. *Nature Methods* **3**, 623–628 (2006).
